# The Tricuspid Valve is Transcriptionally Active During Prolonged Pressure Overload, Right-Sided Heart Failure, and Valve Regurgitation

**DOI:** 10.1101/2025.07.21.666043

**Authors:** Austin Goodyke, Boguslaw Gaweda, Magda Piekarska, Sanjana Arora, Mason Westgate, Renzo Loyaga-Rendon, Milena Jani, Manuel K Rausch, Aitor Aguirre, Jeremy W Prokop, Tomasz A Timek

## Abstract

**Background:** Right-sided heart failure (RHF), in the presence of tricuspid valve regurgitation (TR), can result from left-sided heart failure (LHF), pulmonary hypertension (PH), or heart malformations. The occurrence of RHF and TR represents a critical indicator of hospitalization rates and all-cause mortality. However, RHF has remained understudied, specifically with respect to the tricuspid valve, with few animal models to investigate the transformative processes and identify novel interventions.

**Methods:** Using the outbred sheep (*Ovis aries*) model of pulmonary artery banding (PAB) that induces RHF and TR, we generated three batches of ribosomal reduced RNA sequencing for 354 samples (NCBI SRA PRJNA1182691) containing right ventricle, left ventricle, each tricuspid valve leaflet, each mitral valve leaflet, and the pulmonary artery that represents both male and female sheep. The reads were assembled into a *de novo* sheep heart transcriptome for differential analysis.

**Results:** The *de novo* sheep heart transcriptome enhanced transcript mapping of reads by 43-45% in the heart valves relative to the known sheep reference transcriptome. The identified transcripts produce validated tissue-specific pathways in ventricles (2,756 isoforms), pulmonary arteries (535 isoforms), and valves (1,215 isoforms), with transcript differences between the mitral and tricuspid valve involved in extracellular and endocrine signaling. The transcriptome also produced robust sex differences encoded by sex chromosomes and autosomes, highlighting epigenetic and sex hormone differences in the heart. Echocardiography and differential expression suggest that 8 weeks after PAB, the right ventricle has extensive morphological changes and known stress-induced lipid processing dysregulation. At 16-weeks post-PAB, tricuspid valve leaflets show the most significant transcriptional changes, with alterations in endocrine and immune pathway genes involved in cellular and extracellular remodeling. Genes within the tricuspid valve with differential expression and known human or mouse heart phenotypes include *FLNA*, *LTBP4*, *VDR*, *CR2*, *PIGQ*, *CENPF*, *ACKR3*, *CR1*, *KLF2*, and *HIF3A*.

**Conclusions:** This project highlights the complexity of heart valve tissues and their transcriptional activity in a sheep model of RHF. It suggests potential therapeutic interventions in heart valve remodeling in PAH, RHF, and TR. This work highlights the need for further human and model organism research into the dynamic valve cells and genes.

**Clinical Perspective:** *What Is New?:*

- Improved cardiac and valve specific transcriptome mapping through *de novo* transcriptome for clinically relevant ovine model.
- Tricuspid valves show a sex dependent and active transcriptional response to pulmonary artery banding induced pulmonary hypertension and right sided heart failure.
- Transcriptional phenotypes in ovine model mirror known human heart disease phenotypes.
- Several transcripts identified with therapeutic potential for treating pressure overload conditions such as pulmonary hypertension.

*What Are the Clinical Implications?:*

- Tricuspid valve remodeling due to pulmonary hypertension is accompanied by transcriptional alterations, and mechanical alterations alone may not be sufficient to address valve insufficiency.
- Valve and ventricle sex specific gene expression changes following pulmonary hypertension indicate a potential role for hormonal influences and a need for personalized treatment strategies.
- Improved patient outcomes for right heart failure, including diagnosis, early detection, and improved treatment strategies can be elucidated through the ovine model and supported through expanded interrogation of human tissues.

## Introduction

Heart failure is often the result of diverse pathologies that impact cardiac output, such as hypertension, coronary artery disease, myocardial infarction, and chronic kidney disease. The more than 64 million individuals with heart failure represent a global burden of unfavorable morbidity and mortality, poor quality of life, and billions of dollars in excess clinical costs.^1,2^ Early detection can prevent heart failure progression; however, in cases where medical interventions are limited, the progression of heart failure results in high mortality, and in some patients, worsening heart failure can progress even with interventions.^3^ While many patients suffer left-sided heart failure (LHF), right-sided heart failure (RHF) can signify late-stage disease progression that is highly associated with exercise intolerance and end-organ damage.^4^ While LHF is well studied, there remains a paucity of insights into the molecular mechanisms of RHF.^5,6^

The right ventricle (RV) has a higher capacity than the left for volume expansion but at the cost of thinner ventricular walls, where afterload elevation can be associated with decreased stroke volume and venous blood pooling.^7^ Outside of LHF dysmorphology that alters the RV morphology and pulmonary pressure, RHF is highly associated with pressure changes within the pulmonary artery (pulmonary hypertension-PH, chronic obstructive pulmonary disease, pulmonary embolism), dysmorphology of the tricuspid valve (stenosis, regurgitation), or genetic conditions of cardiomyopathy.^8^ The presence of tricuspid regurgitation (TR), in RHF or PH is associated with elevated hospitalization rates and all-cause mortality.^9^ TR has been deemed a public health crisis, with more than 1.6 million patients in the US being diagnosed with mild or severe TR, which carries a significant correlation with survival.^10,11^ Yet, over 90% of these patients are not offered surgical repair or treatment for TR. There is a critical need for model organisms to define RHF and TR molecular mechanisms that enable preclinical studies for TR intervention within humans.

Sheep (*Ovis aries*) have cardiovascular physiological similarities to humans, including comparable heart size, hemodynamics, and ability to translate clinical methodologies.^12^ The outbred nature of sheep lends a genetic pattern similar to humans, with extensive heterogeneity in the cardiovascular system. The sheep pulmonary artery banding (PAB) model mimics PH, resulting in right ventricular overload, RHF, and TR.^13^ Our team has shown that the PAB model enables extensive dynamic modeling and mechanistic insights of RHF and TR, elucidating the annular dilation with reduced annular area contraction of the RV.^14^ Within PAB, the myocardial stiffness results in ventricular stiffening^15^, while ventricular morphological changes are associated with tricuspid valve (TV) remodeling.^16^ Examination of control and PAB TV leaflets show transcriptional inflammation-related changes, while histology suggests cellularity and extracellular matrix composition changes.^17^ However, the knowledge of gene transcripts within the sheep heart remains a significant challenge in translating the sheep data into humans.

In the current study, we developed 354 gene expression datasets for male and female control or PAB (8-weeks, 12-weeks, or 16-weeks post banding) sheep hearts that include individual leaflets of the tricuspid and mitral valve (MVs) in addition to the RV, left ventricle (LV), and pulmonary artery (PA). This extensive database enables a *de novo* assembly of sheep heart transcripts that significantly enhances mapping and discovery, showing the sheep model’s extensive overlap with human genes associated with heart failure and TR.

## Methods

### Sheep surgery and PAB procedure

Animals were housed and cared for at Michigan State University large animal facility with approval from the Institutional Animal Care and Use Committee, under PROTO202100064 approved 7-15-21. A detailed video of the surgical procedure is available online.^18^ In brief, healthy adult male and female Dorset sheep were anesthetized, intubated, mechanically ventilated, and a limited left thoracotomy was performed. The main PA was then encircled with an umbilical tape proximally to its bifurcation (Figure 1A) and tightened to the brink of hemodynamic stability. Subsequently, three sets of animals were monitored for 8, 12, or 16 weeks. Reversal of the PAB procedure was conducted in a subset of animals in experiment 2 after 8 weeks of PAB and echocardiographic confirmation of severe TR, in which a right thoracotomy was performed to remove the umbilical tape from the pulmonary artery, the animals were then recovered another 8 weeks. At the terminal endpoint, all animals were anesthetized, and epicardial echocardiography was completed to assess biventricular function and valvular competence. Animals were then euthanized through the administration of sodium pentothal, the heart was excised, and samples were collected into RNA-later from the RV, LV, PA (before and after the band), and each leaflet of the tricuspid and MV (Figure 1B). The sheep were divided into three experiments based on the length of PAB; experiment 1 (13 PAB for 16 weeks with 12 paired controls), experiment 2 (7 PAB for 8 weeks, 8 PAB for 8 weeks, debanded and recovered for 8 weeks, and 10 paired controls), and experiment 3 (6 males PAB for 12 weeks, 8 females PAB for 12 weeks) (Figure 1C). All samples were processed in batches for each experiment.

**Figure 1.**
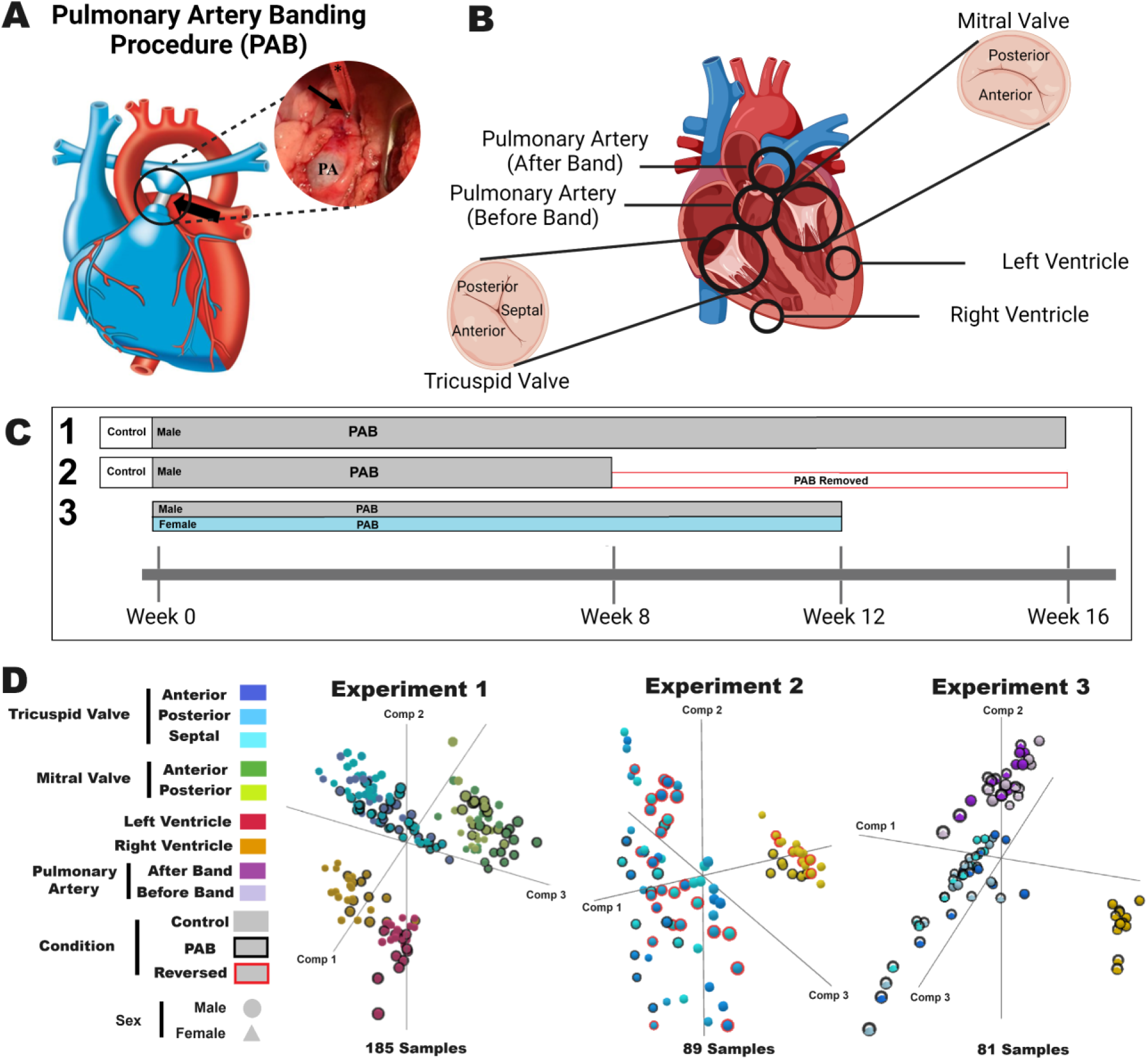
Experimental datasets generated for sheep hearts. **A)** Pictorial of the pulmonary artery banding (PAB) location on the sheep heart. **B)** Location of sample collections from sheep hearts within the current study. Generated with BioRender. **C)** Timeline for sample collections from each of the three experiments. **D)** Principal Component Analysis (PCA) for all 354 samples collected and run for RNA sequencing relative to the reference sheep transcriptome.

### Tissue processing and RNA sequencing

Excised tissue was stored in RNALater (Thermo Fisher) following surgery, incubated at 4°C overnight, and frozen at -80°C. RNA extractions were performed on the tissue using a TRIzol extraction kit (Invitrogen) and an RNeasy MinElute Cleanup kit (Qiagen). Stranded total RNA library preparation with ribosomal reduction was performed with paired-end sequencing at Van Andel Research Institute Genomics Core (Experiment 1) or Azenta Life Sciences (Experiments 2 and 3) to an average depth of around 30 million reads.

### Bioinformatic Analysis

The fastq files were deposited into NCBI SRA under BioProject PRJNA1182691 and processed through FastQC to assess the quality control of each dataset. Reads were aligned to the ARS-UI_Ramb_v3.0 *Ovis aries* transcriptome using Salmon^19^ quasi alignment. Principal Component Analysis (PCA) was performed in NetworkAnalyst^20^, and significant transcripts were identified for transcripts with at least one sample at one transcript per million (TPM) using a Log2 fold change cutoff of 1 and a Bonferroni corrected p-value of <0.05. All sample details and mapping data relative to the reference transcriptome can be found at https://doi.org/10.6084/m9.figshare.27228696.v1. The *de novo* assembly was created using 39 paired-end datasets representing all three experiments, tissues, conditions, and sexes assembled into a Trinity^21^ transcript map annotated for open reading frame protein sequences using Transdecoder and processed for amino acid annotations with Trinotate.^22^ Targeted motif predictions were based on the Eukaryotic Linear Motif tools.^23^ Annotation files for the de novo assembly can be found at https://doi.org/10.6084/m9.figshare.27236733.v1. Sex differences for each tissue were annotated using experiment 3 data with Log2 fold change cutoff at one and a p-value <1E-30. Tissue-specific transcripts were annotated based on Log2 fold change (each tissue relative to all tissue values) cutoff of 2, and Bonferroni corrected p-value of <0.05.

Heatmaps were generated using the Broad Institute Morpheus tools (https://software.broadinstitute.org/morpheus). Using a correlation coefficient, linear regression was performed for various Echo traits relative to the *de novo* transcripts. The *de novo* mapping values are available at https://doi.org/10.6084/m9.figshare.27693090.v1. Genes with significance from experiments 1 or 2 were filtered through the Online Mendelian Inheritance in Man (OMIM)^24^ database for cardiovascular phenotypes, Open Target Platform (OTP)^25^ for TR (HP_0005180) or heart failure (EFO_0003144), the International Mouse Phenotyping Consortium (IMPC) for genes altering heart traits,^26^ and google scholar relative to "heart failure."

## Results

### Echocardiographic changes from PAB

A PAB procedure was performed on sheep to establish PH-induced RHF (Figure 1A). Mitral (anterior-AML and posterior-PML) and tricuspid (anterior-ATL, posterior-PTL, and septal-STL) leaflets, along with left ventricle (LV), right ventricle (RV) and pulmonary artery (PA) were collected from three independent experiments (Figure 1B). These experiments compared PAB (N=13) to control (N=12) following 16 weeks of banding (experiment 1), control (N=10) to PAB following 8 weeks of banding (N=7) or to PAB following 8 weeks of banding (with severe TR) and 8 weeks of recovery (with severe TR) (N=8) (experiment 2) and male (N=6) to female (N=8) following 12 weeks of banding (experiment 3) (Figure 1C).

Clinical assessment of the animals at the end of the three independent experiments demonstrated significant changes associated with PAB (Figure 2). PAB resulted in significantly different pulmonary pressures following 16 weeks of banding, with mean PA pressure (MPAP) increasing from 19±4 to 30±4 mmHg (p<0.001) in experiment 1. Echocardiographic assessment of these animals identified several significantly different metrics indicative of RHF, including increased right atrium area (RAarea) (p<0.001), TR (p<0.001), RV diameter (RVd1) (p<0.001), and tricuspid annular diameter (p<0.001), with decreased tricuspid annular plane systolic excursion (TAPSE) (p<0.001) and RV fractional area contraction (RVFAC) (p<0.001) for animals undergoing the PAB procedure; while left heart assessments remained unchanged.

**Figure 2.**
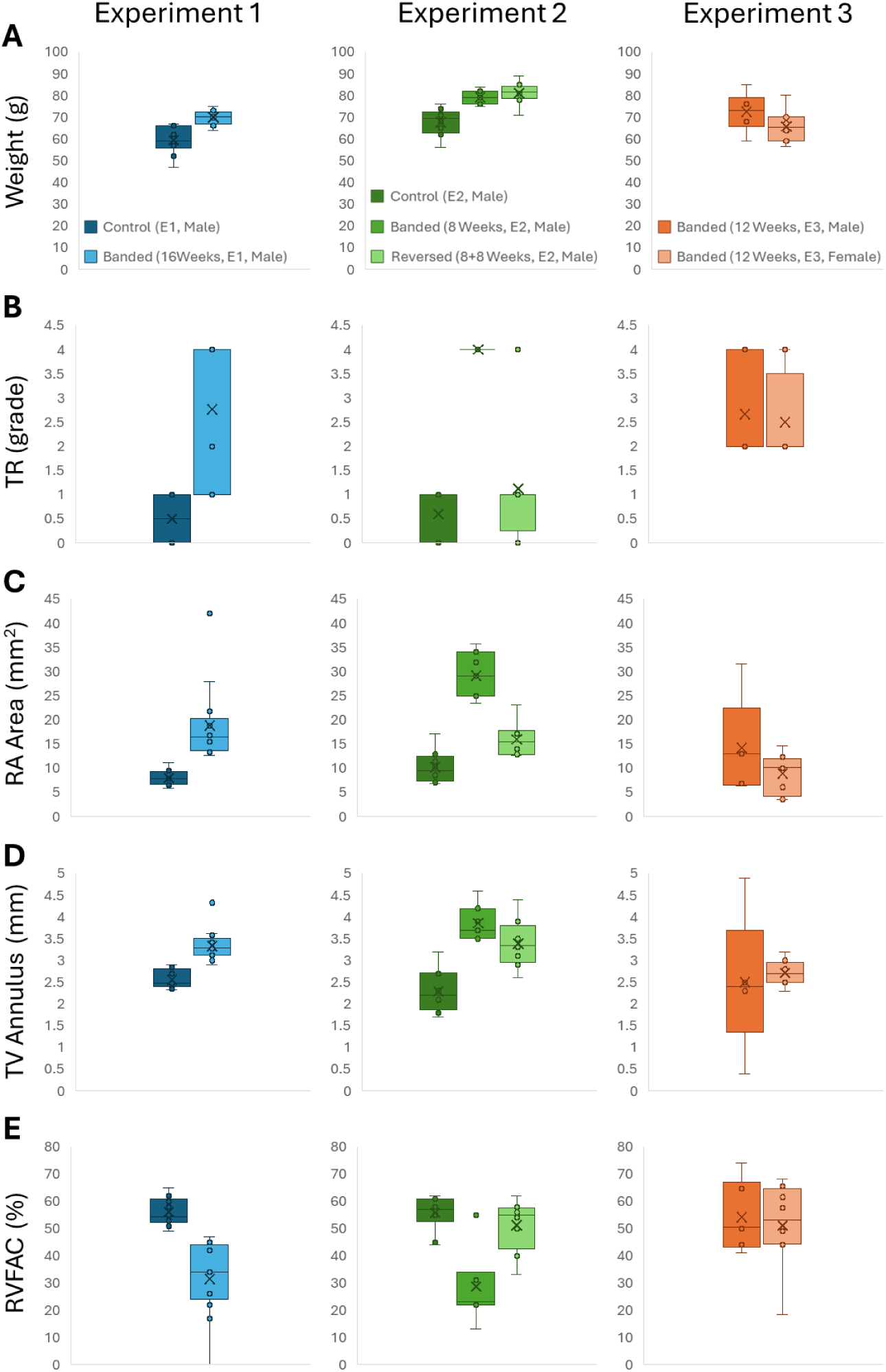
Clinical cardiovascular outcome data for experiments contributing to de novo transcriptome. Data for control and 16-week banded animals from experiment 1; control, 8-week banded, and 8-week banded 8-week reversed animals from experiment 2; as well as male 12-week banded and female 12-week banded animals from experiment 3 including **A)** weights, **B)** tricuspid regurgitation grade, **C)** right atrium area, **D)** tricuspid valve annulus, and **E)** right ventricle fractional area contraction.

Experiment 2 assessed the reversibility of PAB, analyzing the differences between control, severe TR (TR=4) at 8 weeks post-PAB (PAB), and severe TR at 8 weeks with PAB reversed and recovered for 8 weeks (reversed). In comparison to controls, PAB animals had significant increases, similar to those observed in experiment 1, including MPAP (p<0.001), TR (p<0.001), RAarea (p<0.001), RVd1 (p<0.001), and TAPSE (p=0.004). Reversal of the PAB procedure resulted in many clinical metrics returning to normal, with few metrics being significantly different from control except for TV AD (p=0.020). At the same time, the reversed group significantly differed from PAB animals in TR (p<0.001), RAarea (p<0.001), RVd1 (p<0.001), RVFAC (p=0.002), and TAPSE (p=0.033). Experiment 3 compared PAB male and female sheep, which responded similarly to the PAB procedure with no significant differences observed in clinical metrics.

### RNA sequencing mapped to the reference sheep transcriptome

RNA sequencing produced high-quality data for all experiments and each tissue cluster based on PCA (Figure 1D). Control and PAB for each tissue and different leaflets within each valve clustered together, indicating that tissue specificity was the most significant factor in all three components of the PCA.

We identified significant transcripts for statistical comparisons over all three experiments using a Bonferroni corrected p-value relative to the number of transcripts with at least one sample greater than 1 TPM (Figure 3). In experiment 1, ignoring the vast differences between valves and ventricles, we identified the most significant tissue differences between the MV and the TV (154 transcripts), followed by the LV to RV (65 transcripts). In the PAB animals, the number of significant transcripts of the MV to TV comparisons was elevated to 826, indicating the valve undergoes significant alteration in expression. Comparing control to PAB sheep for each tissue elucidated the most significant changes in the TV (236 transcripts). This suggests that at 16 weeks post PAB of experiment 1, the TV remains transcriptionally altered more than the RV.

**Figure 3.**
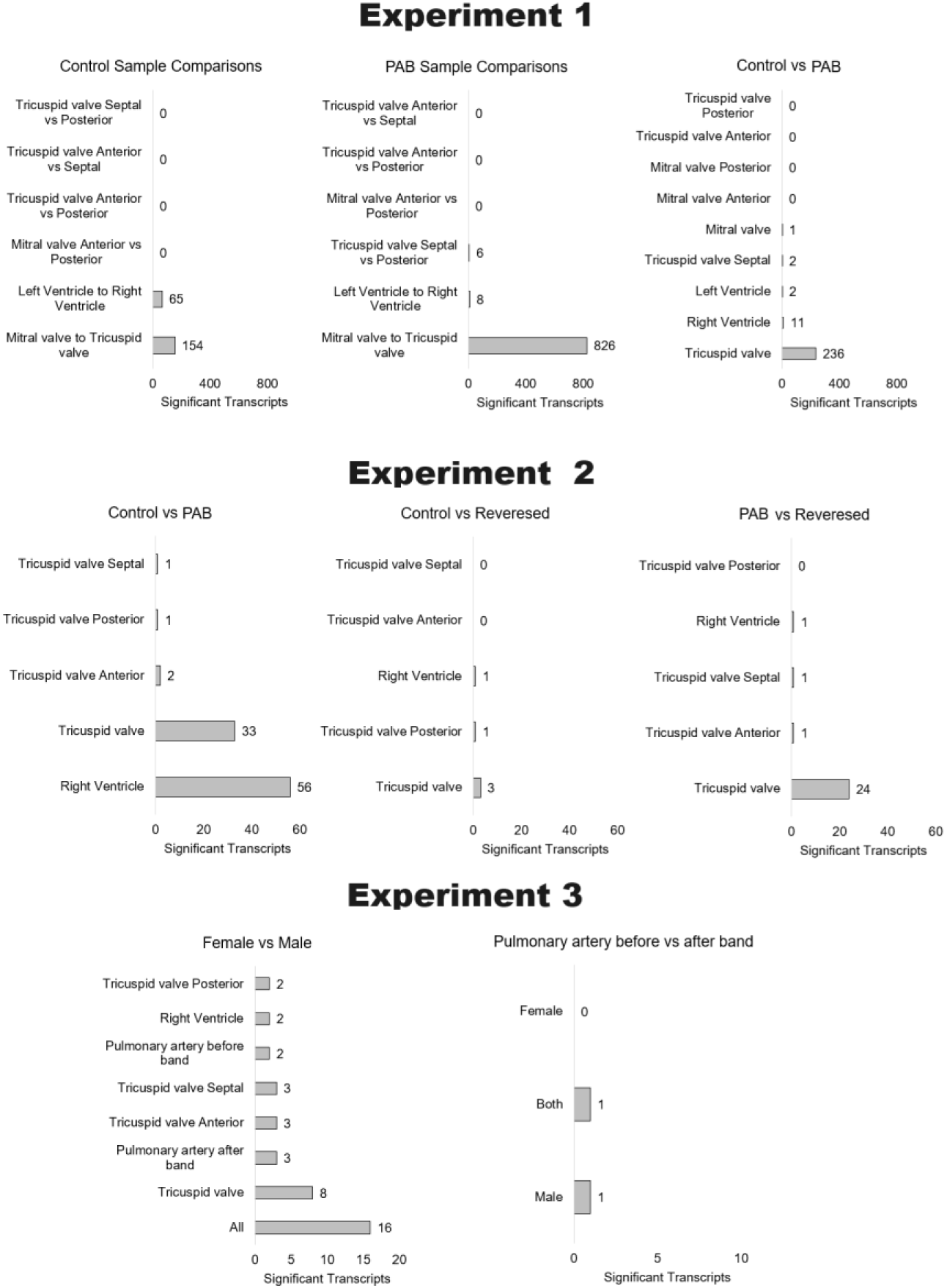
**Statistical comparisons of reference alignment statistics over the three experiments.**

Within experiment 2, we observed the most significant differences in control vs. 8 weeks post PAB for the RV (56 transcripts), with fewer changes observed within the TV (33 transcripts). The control and reversed PAB animals had few significant transcripts detected. The 8-week post-PAB animals relative to animals that received 8 weeks PAB and 8 weeks recovery from the band removed (reversal group) had the most significant transcript difference observed within the TV (24 transcripts), suggesting that reduction of pressure within the right side of the heart significantly changes the TV more so than other tissues. The combination of experiments 1 and 2 suggests that the TV is the most transcriptionally active region within the heart in PAB.

As experiments 1 and 2 were performed in all male sheep, we added experiment 3 to address sex differences by studying a cohort of male and female sheep 12 weeks post-PAB. These animals show significant sex differences across all animals. However, the number of identified transcripts was relatively low, suggesting a poor representation of sex-specific transcripts within the reference transcriptome. We additionally included RNA sequencing of the PA at locations flanking the band of the animals, where mapping of reads was relatively low, and few significant transcripts were detected relative to the side of the band.

### Building a de novo Sheep heart transcriptome

Addressing the percentage of alignment of reads to the reference transcriptome for valve and PA samples suggested sub-optimal alignments (Figure 4A, x-axis) for performing downstream analysis such as gene ontology. This indicates that the sheep heart valves and arteries are poorly defined in the reference transcriptome assembly relative to the heart ventricles. To correct this mapping issue, we built a *de novo* assembly of the reads into transcripts. As the number of sample inputs is limited by computing RAM, we selected 39 of the samples for processing with 1.5TBs of RAM, which included a representative tissue sample of each experiment for each conditional group, and experiment 3 also included one sample of each for either male or female. Quasi-alignment of the reads for all samples relative to this *de novo* assembly significantly enhances mapping to an average of 76.22±5.47% relative to the 38.79±11.9% for the reference transcriptome, a 37.43% improvement. Focusing on the MV (35.87±1.63 vs. 81.32±0.91%), TV (31.56±4.75 vs. 74.58±4.84), and PA (37.28±2.12% vs.

**Figure 4.**
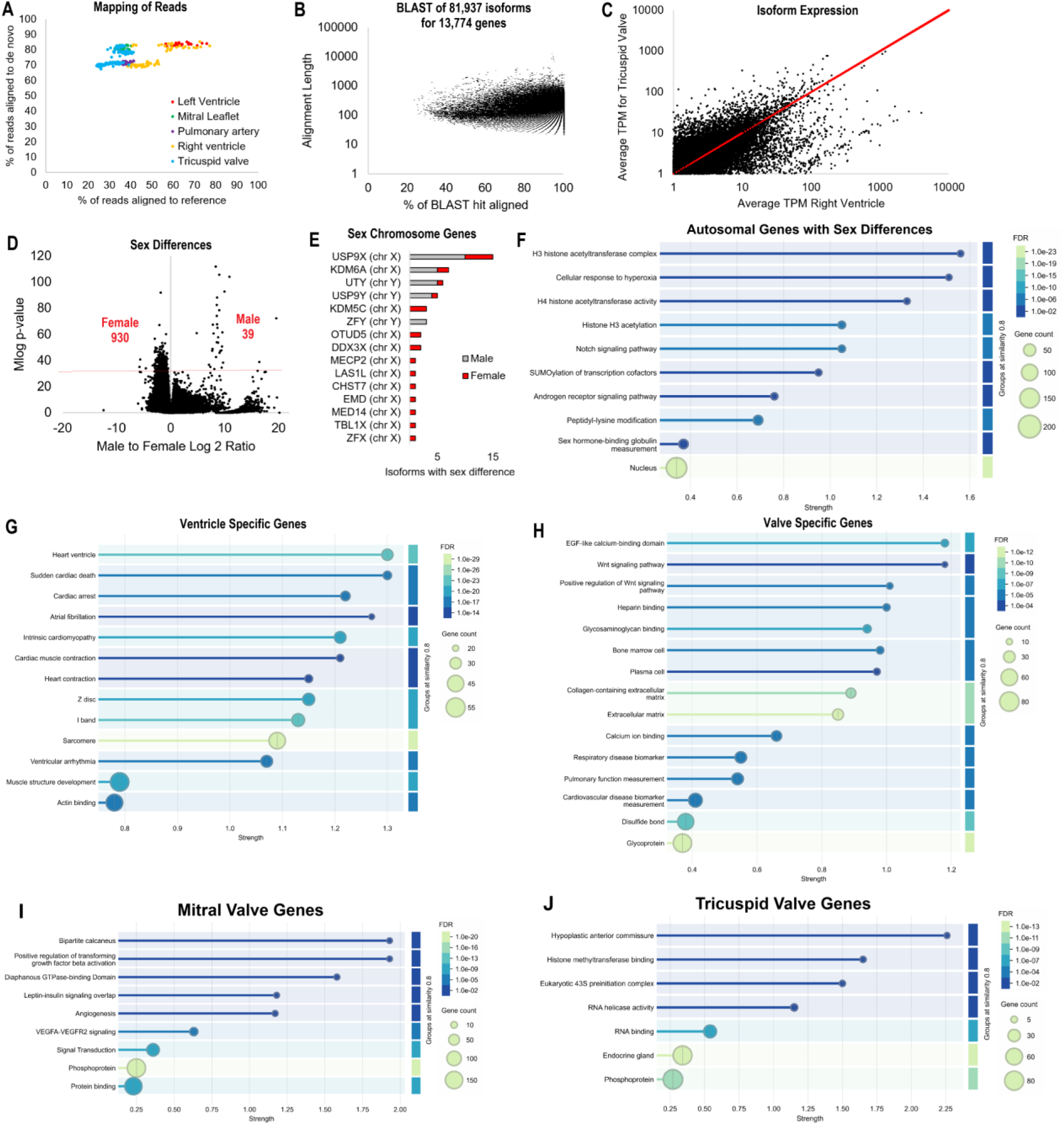
Mapping of reads to a novel *de novo* sheep heart transcriptome. **A)** The mapping of each dataset to the reference sheep transcriptome (x-axis) relative to the *de novo* transcriptome (y-axis) for each labeled tissue. **B)** The BLAST analysis of transdecoder identified protein sequences within the *de novo* assembly for the percent similarity to BLAST hit (x-axis) relative to the portion of the fragment identified in amino acids (y-axis). **C)** The average expression of each transcript from panel B in the right ventricle (x-axis) relative to the tricuspid valve (y-axis). **D)** Transcripts with a significant difference between male and female samples. **E)** Transcripts from panel D that match the sex chromosomes (chr X and chr Y). **F-J)** Gene ontology enrichment for autosomal genes enriched in sex differences (panel **F**), transcripts significantly elevated in the left and right ventricles (panel **G**), transcripts significantly elevated in the mitral and tricuspid valves (panel **H**), or transcripts elevated in control mitral (panel **I**) vs. tricuspid (panel **J**) valves.

71.78±0.71%) we see an enhancement of 45.4%, 43.0%, and 34.5% (Figure 4A). This marked improvement represents a significant need for studying tissues such as heart valves in model organisms, representing the underdeveloped knowledge of gene expression for these heart regions.

Assembled transcripts were assessed for an open reading frame that codes for proteins that match a known protein based on BLAST analysis (Figure 4B). A total of 81,937 transcripts from 13,774 unique proteins were identified with a range of similarities to reference within BLAST and amino acid length. Some of these transcripts were present in all tissues, while some showed expression bias in tissues such as the RV or TV (Figure 4C).

One of the most significant improvements for the *de novo* assembly relative to the reference alignments was identifying sex differences within the sheep heart. A rigorous p-value cutoff of <1x10^-30^ produced 39 transcripts elevated in the male hearts and 930 within female hearts (Figure 3D). Of these sex differences, multiple transcripts map to genes on the sex chromosomes, including chromosome X genes *USP9X*, *KDM6A*, *KDM5C*, *ZFX*, *OTUD5*, *DDX3X*, *MECP2*, *LAS1L*, *CHST7*, *EMD*, *MED14*, *TBL1X* and chromosome Y genes *UTY*, *USP9Y*, *KDM5D*, *ZFY* (Figure 4E). Genes with sex differences coded from the autosomes are enriched for pathways including histone epigenetic regulations, notch signaling, SUMOylation, androgen receptor, and sex-hormone-binding globulin measurements (Figure 4F).

Addressing tissue specificity, we observed six transcripts that were enriched in the LV (within the *PEBP4*, *LMOD2*, *ATG9A*, *POL*, *TNNT2*, *IASPP* genes), 64 in the RV, 4 in the TV (*PDILT*, *CO6A3*, *LGR6*, *EXT1*), and 535 in the PA. The RV was enriched for genes involved in sarcomere organization (GO:0045214, enrichment strength 2.08, false discovery rate-FDR 3e-07), myofibril (GO:0030016, enrichment strength 1.62, FDR 2.9e-15), and cardiomyopathy (DOID:0050700, enrichment strength 1.53, FDR 6.7e-06). Of the 535 transcripts over 213 genes enriched within the PA, pathway enrichments include smooth muscle cells (PMID:26241044, enrichment strength 1.85, FDR 6.1e-08) and actin filament bundle (GO:0032432, enrichment strength 1.15, FDR 2.6e-05).

Many protein-coding genes were enriched in the ventricles (shared between right and left) or the valves (tricuspid and mitral). The ventricles had 2,756 elevated transcripts from 1,166 genes enriched for terms including heart ventricle, cardiac arrest, cardiomyopathy, and heart contractions (Figure 4G). The top genes include Titan (65 transcripts), Ryanodine receptor 2 (32 transcripts), Myomegalin (24 transcripts), and Obscurin (23 transcripts), which are all well-documented as muscle-elevated proteins.

The valves were enriched for 1,215 transcripts from 540 genes enriched for terms including Wnt signaling, heparin-binding, bone marrow cells, extracellular matrix, and respiratory/pulmonary function (Figure 4H). The top genes include LINE-1 retrotransposable element ORF2 protein (32 transcripts), Fibrocystin-L (30 transcripts), Hemicentin-1 (20 transcripts), and Cell surface hyaluronidase CEMIP2 (18 transcripts).

A total of 397 transcripts in the PA, 45 in the valves, and 5 in the ventricles have tissue specificity and sex differences with the top genes, including Vimentin (valve elevated with male bias), Insulin-like growth factor II (valve elevated with male bias), Lumican (valve elevated with male bias), Myosin-11 (PA with female bias), and Filamin-A (PA with female bias). Overall, the tissue enrichment of transcripts properly reflects the biological pathways of the heart, validating the *de novo* transcript maps as biologically meaningful.

The TV and MV also showed significant differences in control sheep with the *de novo* assembly, similar to the reference alignment. There were 477 transcripts from 412 genes significantly elevated in the MV. The MV is enriched for genes involved in various extracellular signaling pathways (TGF-beta, leptin, insulin, VEGFA) and signal transduction (Figure 4I). There were 197 transcripts from 188 genes significantly elevated in the TV. The TV is enriched for genes involved in RNA biology and endocrine function (Figure 4J).

Our novel sheep heart *de novo* transcriptome represents a significant enhancement into sex differences and regions of the heart. With the enhanced transcript maps, we recalculated significant transcripts in PAB of experiment 1 and PAB or PAB reversal of experiment 2. There were 192 transcripts from 159 unique genes powered for significant difference with Bonferroni correction in any of the comparisons, as detailed below.

Four transcripts were significant over all tissues of experiment 1 for control vs. PAB, including two transcripts for the *RAB1A* gene, one in the *AMD* gene, and one in the *FCHO2* gene. A 198 amino acid coding transcript for *RAB1A* (TRINITY_DN205354_c1_g2_i8 of our *de novo* assembly), a small GTPase involved in vesicle trafficking, is found at 2.69±0.14 in control and 0.61±0.11 in PAB samples with a Log2 fold change of -2.15 and a p-value of 6.1e-24. The protein is 100% homologous to the human RAB1A protein (UniProt P62820). A 228 amino acid coding transcript (TRINITY_DN210291_c2_g1_i6) that shares 132 amino acid homology to a known bovine *AMD* gene is found in control tissues at 4.27±0.37 and PAB tissues at 0.48±0.17 with a Log2 fold change of -3.14 and a p-value of 3.90e-16. The coded protein has no homology to humans, no strong annotated domains based on Pfam, a disordered sequence based on InterPro, and multiple predicted linear motifs of extracellular processing based on ELM analysis. Finally, a 109 amino acid coding transcript (TRINITY_DN169630_c0_g1_i2) with homology to the F-BAR domain only protein 2 is expressed in control tissues at 0.25±0.7 and PAB tissues at 1.10±0.12 with a Log2 fold change of 2.16 and a p-value of 3.31e-09. The protein is 95% similar to the human FCHO2 for amino acids 728-810 (UniProt Q0JRZ9) and includes a muniscin C-terminal mu homology domain known to alter muniscin localization in endocytosis. These three protein-coding genes represent an interesting observation of changes throughout the RV, LV, TV, and MV due to PAB, suggesting global alterations in the sheep heart model.

Within the RV of experiment 1, 10 transcripts (from 9 genes) are significantly higher in control vs PAB, and 17 transcripts (from 14 genes) are higher in the PAB animals. The genes have no pathway enrichments. Eight transcripts were from the ventricle elevated list of Figure 4G, and none had sex differences. The most significant transcript codes for a 383 amino acid protein with 90% homology to the human Hydroxysteroid dehydrogenase-like protein 2 (*HSDL2*, a ventricle enriched gene) with an average expression in the RV control samples of 11.08±0.46 and PAB samples of 3.91±0.39 with a Log2 fold change of -1.50 and a p-value of 4.14e-11. The protein has predicted HSDL2_SDR_c and SCP2 InterPro domains that maintain its lipid metabolism functions. The highest expression levels of significant genes included a 220 amino acid coding transcript with 92% homology to the human Ferritin heavy chain (*FTH1*, control 242.33±12.46, PAB 564.89±50.81, p-value 2.58e-07) and 660 amino acid coding transcript with 93% homology to the human Nebulin-related-anchoring protein (*NRAP*, control 37.12±3.20, PAB 95.23±8.37, p-value 1.82e-07).

Within the TV of experiment 1, 91 transcripts (from 75 genes) are significantly higher in control vs. PAB, and 8 transcripts (from 8 genes) are higher in the PAB animals. The genes are enriched for EGF-like domain (IPR000742, enrichment strength 1.22, FDR 3.57e-06), serine-type endopeptidase activity (GO:1900005, enrichment strength 2.42, FDR 4.13e-02), integrin-medicated signaling (GO:2001046, enrichment strength 1.96, FDR 1.60e-02), low-density lipoprotein particle binding (GO:0030169, enrichment strength 1.97, FDR 1.00e-03), transforming growth factor beta binding (GO:0050431, enrichment strength 1.69, FDR 2.34e-02), and protein complex involved in cell-matrix adhesion (GOCC:0098637, enrichment strength 1.71, FDR 1.46e-02). The most significant isoform coded for a 236 amino acid protein with 39% homology to the human CD209 antigen (*CD209*, control 3.25±0.67, PAB 1.25±0.08, p-value 3.79e-16). The highest expressed transcript with significance coded for a 429 amino acid protein with no homology to humans but 98.8% homology to the bovine Gelsolin (*GSN*, control 101.73±6.74, PAB 47.86±2.72, p-value 6.91413E-10). The second highest expressed transcript with significance coded for a 333 amino acid protein with 94% homology to the latent transforming growth factor beta binding protein 4 (*LTBP4*, control 26.32±1.31, PAB 13.11±0.90, p-value 5.54e-12).

Within experiment 2, we identified six transcripts in the RV (*ERBIN*, *HS2ST*, *KLHL4*, *CSDE1*, *NHLC2*, *AASS*) and 59 (from 51 genes) in the TV with PAB or reversal of PAB comparisons. The most significant transcript for control vs PAB was a 365 amino acid coding transcript with 95.6% homology to human isthmin-1 precursor (*ISM1*, control 0.48±0.05, PAB 1.32±0.09, PAB-reversal 0.71±0.08, p-value 9.18e-11). The most significant transcript for PAB vs PAB-reversal was a 717 amino acid coding transcript with 81% homology to the human macrophage-expressed gene 1 protein precursor (*MPEG1*, control 3.50±0.20, PAB 5.85±0.29, PAB-reversal 2.87±0.11, p-value 5.65e-11). The most significant transcript for control vs. PAB-reversal was a 275 amino acid coding transcript with 97% homology to the human notch receptor 2 (*NOTCH2*, control 0.41±0.12, PAB 1.89±0.30, PAB-reversal 2.63±0.25, p-value 3.76E-11).

Integrating tissue-specific, sex-specific, and significant transcripts into Spearman’s rank clustering manifests some batch effects (Figure 5A). However, many of these top markers overlap all three experiments for tissue-specificity and, therefore, validate the substantial resources of the novel *de novo* transcript map. Further assessment of the TV significant genes (control vs. PAB) of experiment 1 shows some trends across all valve leaflets, including the MV, with the TV septal leaflet showing the most pronounced differences for these genes in control vs PAB (Figure 5B). In experiment 2, all leaflets show similar changes for the significant genes identified in PAB or PAB-reversal (Figure 5C).

**Figure 5.**
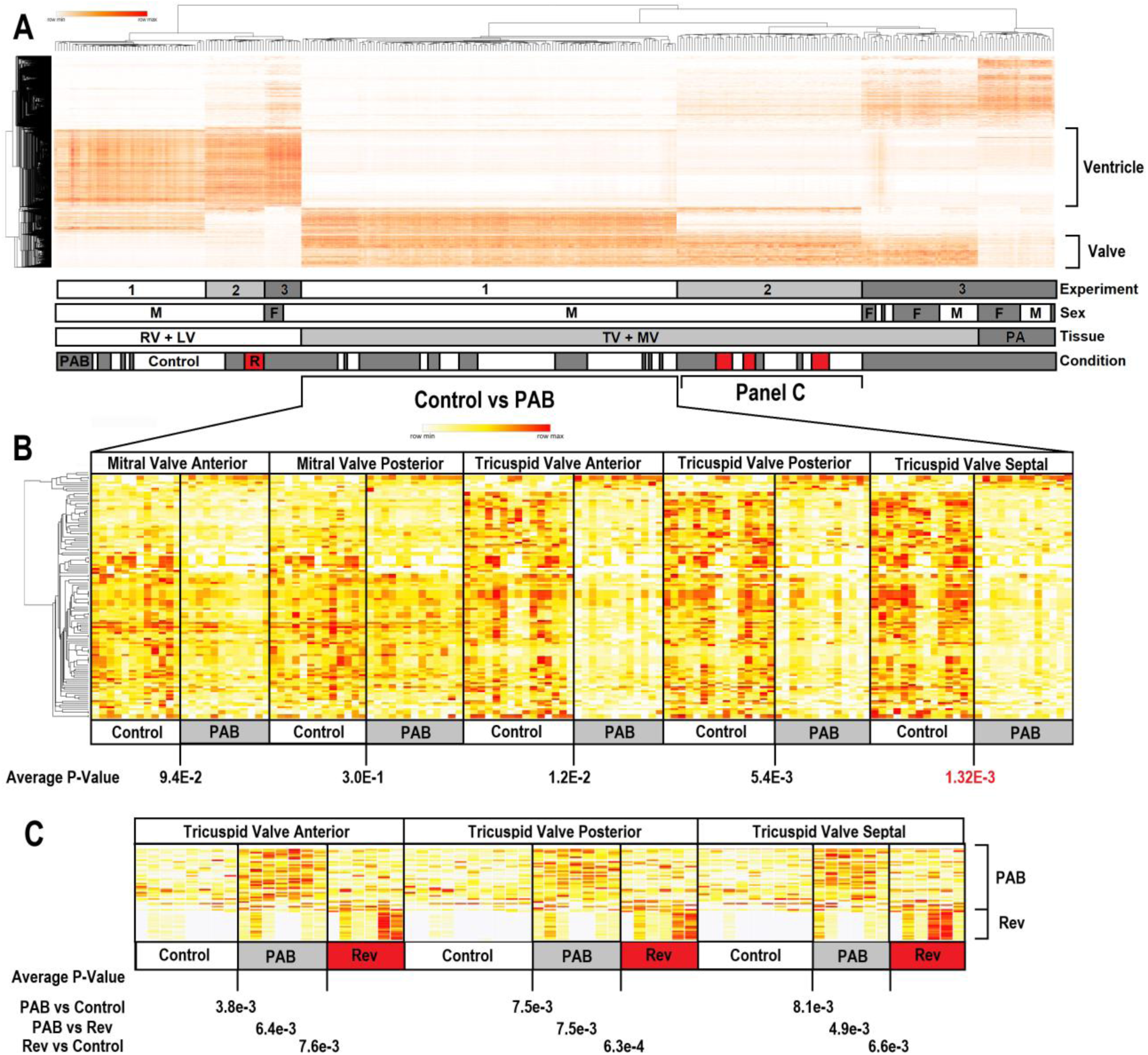
Sample clustering relative to top identified transcripts. **A)** All tissues, conditions, and experiments listed for transcripts with sex differences, tissue specificity, or significant difference of experiment 1 or 2 clustered based on one minus Spearman’s rank as shown on the dendrograms on the top and side. Shown below is a pictorial of the sample labels (M-male, F-female, RV-right ventricle, LV-left ventricle, TV-tricuspid valve, MV-mitral valve, PA-pulmonary artery, PAB-pulmonary artery banding, R-reversed banding). **B)** Expression of the significant transcripts from experiment 1 PAB vs control shown for each leaflet of the mitral and tricuspid valves. At the bottom is the average P-value over all the transcripts for each of the leaflets. **C)** Expression of significant transcripts from experiment 2 control, PAB, and PAB reversal for each leaflet of the tricuspid valves. At the bottom is the average P-value for significant transcripts of each comparison over all the transcripts for each of the leaflets.

### Translating sheep transcripts to heart phenotypes

All significant transcripts from experiments 1 and 2 were screened through various phenotypic datasets to link them to previous insights of heart phenotypes. We identified 23 genes with known human or mouse heart phenotypes (Table 1). Amongst these genes are three transcripts of *FLNA* and 5 in the *CR2* gene with significant differences at 16 weeks PAB in all TV tissues but without significant differences at week 8 PAB or in the PAB reversal (Figure 6), suggesting that pressure overload normalized the expression. The ventricle-specific gene *CD36* showed lower expression in the LV and RV during both 8- and 16-week PAB, which trended towards control levels when PAB was reversed. The phenotypic overlaps of PAB-altered transcripts with known mice and human phenotypes suggest that the sheep model significantly overlaps human physiology.

**Figure 6.**
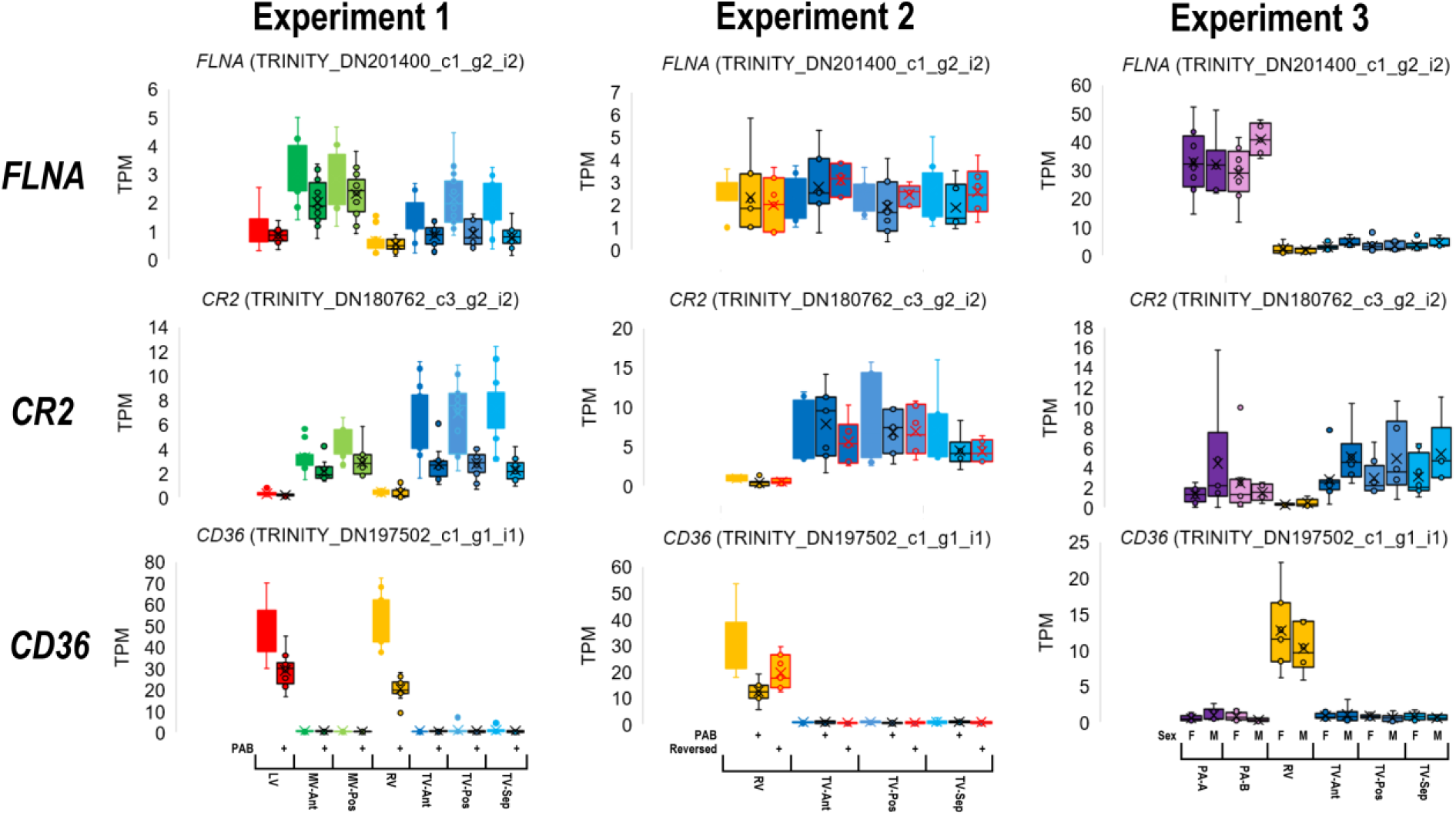
Representative four gene transcripts that showed significant differences between PAB and control sheep. Each experiment is plotted separately to account for batch effect and the group labels are shown at the bottom of the CD36 plots. Color coding is similar to Figure 1 D.

**Table 1.**
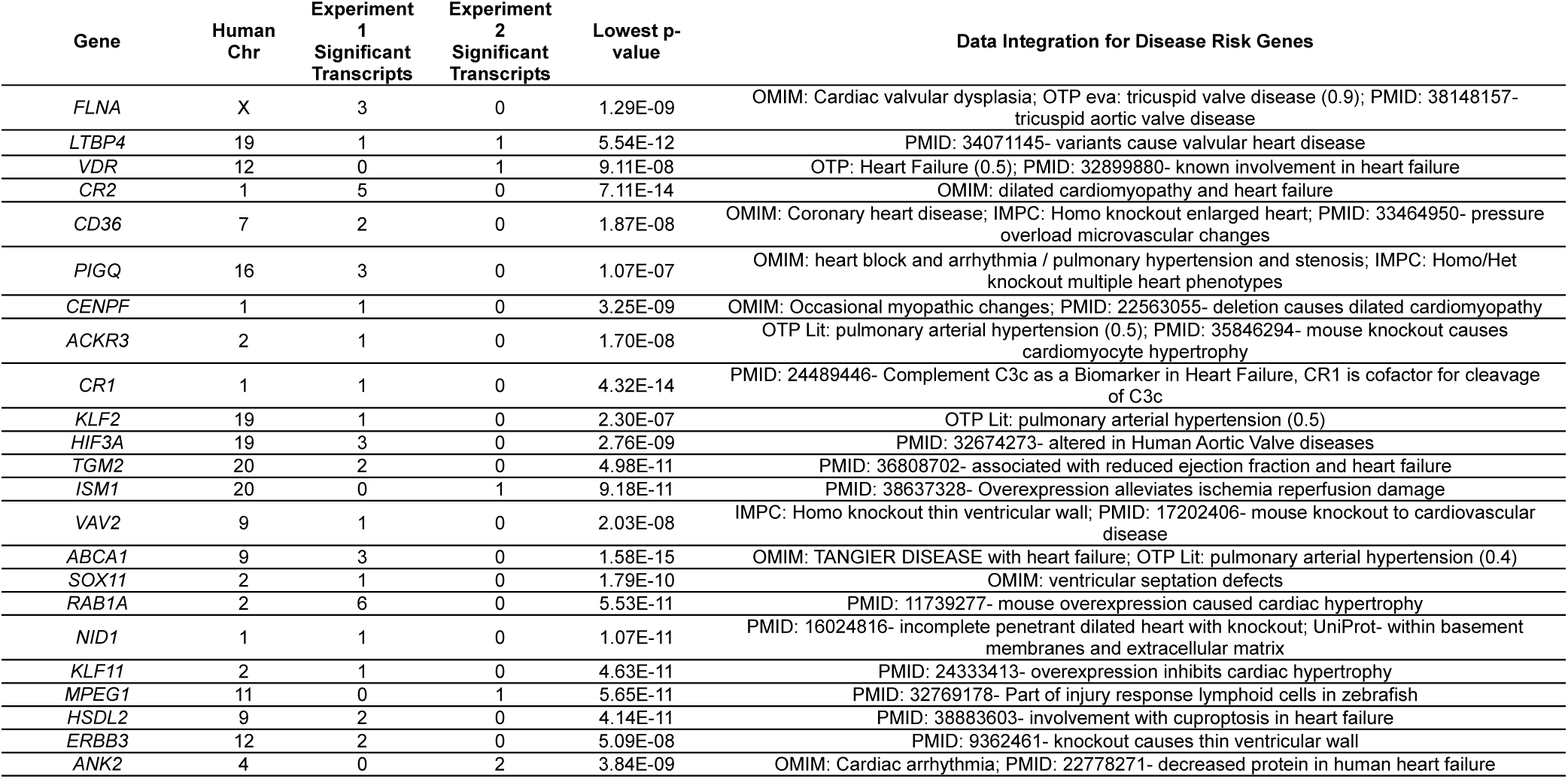
Top genes and transcripts mapped and known disease insights from human. **Footnotes:** OMIM-Online Mendelian Inheritance in Man, OTP-Open Targets Platform, eva-ClinVar based enrichment, PMID-PubMed Index, IMPC-International Mouse Phenotyping Consortium, Lit-literature.

| Gene | Human Chr | Experiment 1 Significant Transcripts | Experiment 2 Significant Transcripts | Lowest p-value | Data Integration for Disease Risk Genes |
| --- | --- | --- | --- | --- | --- |
| FLNA | X | 3 | 0 | 1.29E-09 | OMIM: Cardiac valvular dysplasia; OTP eva: tricuspid valve disease (0.9); PMID: 38148157- tricuspid aortic valve disease |
| LTBP4 | 19 | 1 | 1 | 5.54E-12 | PMID: 34071145- variants cause valvular heart disease |
| VDR | 12 | 0 | 1 | 9.11E-08 | OTP: Heart Failure (0.5); PMID: 32899880- known involvement in heart failure |
| CR2 | 1 | 5 | 0 | 7.11E-14 | OMIM: dilated cardiomyopathy and heart failure |
| CD36 | 7 | 2 | 0 | 1.87E-08 | OMIM: Coronary heart disease; IMPC: Homo knockout enlarged heart; PMID: 33464950- pressure overload microvascular changes |
| PIGQ | 16 | 3 | 0 | 1.07E-07 | OMIM: heart block and arrhythmia / pulmonary hypertension and stenosis; IMPC: Homo/Het knockout multiple heart phenotypes |
| CENPF | 1 | 1 | 0 | 3.25E-09 | OMIM: Occasional myopathic changes; PMID: 22563055- deletion causes dilated cardiomyopathy |
| ACKR3 | 2 | 1 | 0 | 1.70E-08 | OTP Lit: pulmonary arterial hypertension (0.5); PMID: 35846294- mouse knockout causes cardiomyocyte hypertrophy |
| CR1 | 1 | 1 | 0 | 4.32E-14 | PMID: 24489446- Complement C3c as a Biomarker in Heart Failure, CR1 is cofactor for cleavage of C3c |
| KLF2 | 19 | 1 | 0 | 2.30E-07 | OTP Lit: pulmonary arterial hypertension (0.5) |
| HIF3A | 19 | 3 | 0 | 2.76E-09 | PMID: 32674273- altered in Human Aortic Valve diseases |
| TGM2 | 20 | 2 | 0 | 4.98E-11 | PMID: 36808702- associated with reduced ejection fraction and heart failure |
| ISM1 | 20 | 0 | 1 | 9.18E-11 | PMID: 38637328- Overexpression alleviates ischemia reperfusion damage |
| VAV2 | 9 | 1 | 0 | 2.03E-08 | IMPC: Homo knockout thin ventricular wall; PMID: 17202406- mouse knockout to cardiovascular disease |
| ABCA1 | 9 | 3 | 0 | 1.58E-15 | OMIM: TANGIER DISEASE with heart failure; OTP Lit: pulmonary arterial hypertension (0.4) |
| SOX11 | 2 | 1 | 0 | 1.79E-10 | OMIM: ventricular septation defects |
| RAB1A | 2 | 6 | 0 | 5.53E-11 | PMID: 11739277- mouse overexpression caused cardiac hypertrophy |
| NID1 | 1 | 1 | 0 | 1.07E-11 | PMID: 16024816- incomplete penetrant dilated heart with knockout; UniProt- within basement membranes and extracellular matrix |
| KLF11 | 2 | 1 | 0 | 4.63E-11 | PMID: 24333413- overexpression inhibits cardiac hypertrophy |
| MPEG1 | 11 | 0 | 1 | 5.65E-11 | PMID: 32769178- Part of injury response lymphoid cells in zebrafish |
| HSDL2 | 9 | 2 | 0 | 4.14E-11 | PMID: 38883603- involvement with cuproptosis in heart failure |
| ERBB3 | 12 | 2 | 0 | 5.09E-08 | PMID: 9362461- knockout causes thin ventricular wall |
| ANK2 | 4 | 0 | 2 | 3.84E-09 | OMIM: Cardiac arrhythmia; PMID: 22778271- decreased protein in human heart failure |

### Modeling heterogeneous outcomes in an outbred model

The heterogeneous response due to diverse genomic backgrounds and environmental factors is a hallmark of human heart physiology. Given that sheep are an outbred model, and we had relatively high sample sizes, we addressed whether any gene expression may correlate with the levels of heart changes. Echocardiographic assessments of animals from experiment 1 were correlated to significantly differentially expressed isoforms identified by the *de novo* assembly. We observed that more differentially expressed isoforms had a correlation coefficient >0.9 or <-0.9 for RAv (97) than RVEF (4) or TR (2) (Figure 6). Assessment of RAv, RAarea, RVEF, TA, TAPSE, and RVFAC had the most isoforms identified within the RV, while TR assessment had the most in the STL (Figure 7A). Specific isoforms correlating with RAv (*UB2Q2*, *TEC*, *ARID2*), TR (*THL*), and RAv (*YTDC2*) had a positive correlation, while those correlating with RVFAC (*UB2Q2*, *TEC*, *ARID2*), TA (*IDHC*), RVEF (*MIER3*) had negative correlations (Figure 7B). The linear regression suggests that genomic background of the sheep may contribute to the extent of physiological changes within the sheep heart following PAB.

**Figure 7.**
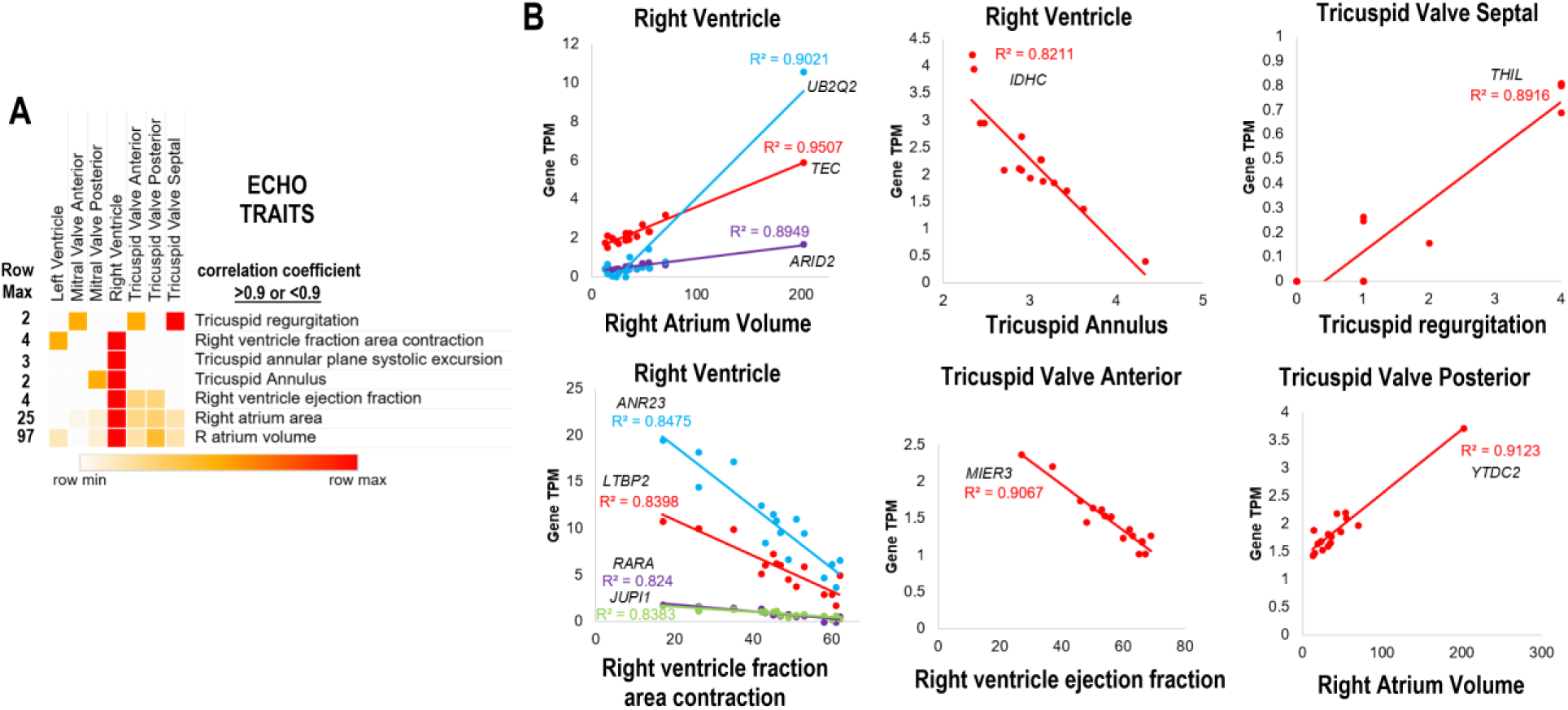
Correlations of gene expression relative to echocardiography (Echo) measured traits. **A)** The number of transcripts from the *de novo* assembly with a correlation coefficient >0.9 or <-0.9 in each tissue RNA sequencing to various Echo measurements. The max number of transcripts in any one tissue is listed to the left. B) Representative gene transcripts shown for various traits and tissues with significant R-squared levels.

## Discussion

Understanding the complex process of RHF and FTR requires model systems, as generating insights from this heterogeneous human disease has proven challenging. The ovine model represents an ideal system for cardiovascular disease as sheep and human hearts are similar on the cellular and molecular level and possess similar heart rates, pressures, and flow dynamics.^27,28^ Sheep model systems also employ an outbred genetic background that explores the heterogenous human RHF physiology missed in inbred small animal model systems that can show strain-specific outcomes.^29^ While this heterogeneous nature poses some intrinsic challenges that require increased sample size, it offers a model system much closer to what is encountered clinically. The current study generated 354 RNAseq datasets publicly shared (NCBI SRA PRJNA1182691) over three divergent experiments to develop a deeply informed heart transcriptome of a model organism for RHF.

The TV transcriptome is poorly defined in healthy and disease states for human and model organisms. Our sheep model provides ample tissue to isolate and compare the expression of each leaflet of the TV relative to the MV. As of early 2025, only 253 runs of "Tricuspid Valve" for RNA have been deposited into NCBI SRA, the primary repository for sequencing experiments. Only fifteen samples of human RNA have been performed for the TV, a remarkably low number. This study represents the most extensive sequencing of RNA within the TV for both control and RHF of any species, with 187 RNA samples of the TV generated in our work representing 74% (187/253) of the current TV expression knowledge.

The TV leaflets maintain similar expression profiles but have significant differences relative to the MV, more so than the RV vs. LV. While anatomically, the differences in MV and TV are well documented,^30^ very few studies have addressed the expression differences. When we first attempted to use the reference alignment, it became clear that our knowledge of valve transcripts was poorly defined based on the percent of read alignment to the reference, thus informing us to develop a *de novo* assembly, which resulted in ∼44% increase in read alignment for both MV and TV. The novel *de novo* transcriptome showed extensive valve enrichment of genes involved in developmental signaling, such as Wnt, with significant differences between mitral and tricuspid, highlighting endocrine signaling differences in line with knowledge of heart developmental pathways.^31^

The PAB model provides a controlled method to induce right ventricular remodeling similar to PH-induced RHF.^32^ Yet, less work has been done on how the TV is altered during RHF. There is little insight into the early transformations of the human heart during PH, of which our PAB model lends insights. Figure 2 shows that in the first 8 weeks following PAB, extensive changes to the RV begin to normalize by week 12 and 16 post-PAB. Our transcriptomic data suggests that 8 weeks post PAB, the RV shows significant changes, while at 16 weeks, the TV is the most altered, which is reversible in controlling TV function with the removal of PAB at 8 weeks. This suggests that in the early phases of RHF, the cardiomyocytes have an early response followed

by delayed valve reconstruction. This is similar to the known MV changes in myocardial infarction (MI) that show passive alterations during the first 8 weeks of damage during MI-induced mitral regurgitation.^33^ Cardiomyocyte-enriched genes such as *LDLR* and *CD36* had decreased expression starting at 8 weeks post-PAB, where it is established as a regulator of fatty acid uptake that protects cardiomyocytes during hypoxic stress such as ischemia.^34,35^ This hypoxic insult is supported by other genes, such as *HIF3A and ISM1* in Table 1.

Our current study supports a delayed active remodeling of valve tissue, which we suggest is driven by endocrine, metabolic, and immune pathways. Under normal physiological conditions, the growth of the valve is regulated by TGF-ß^36^ through valve interstitial cell-extracellular matrix remodeling.^37^ Latent TGF-ß binding protein 4 (LTBP4), a protein with a known protective role in hypoxic-induced tissue damage and valve disorders,^38,39^ was significantly altered in PAB at 8 and 16 weeks, supporting the possible role of TGF-ß in early signaling for valve remodeling. Genes in Table 1, such as Vitamin D receptor (*VDR*),^40^ Atypical chemokine receptor 3 (*ACKR3*), VAV2, *RAB1A, ERBB3,* and *ANK2* support connections of endocrine and signaling circuits to involvement in valve remodeling. KLF transcription factor 2 (*KLF2*) represents a convergence point of endocrine signaling and hemodynamic forces to transcriptional changes within the TV.^41^ Transglutaminase 2 (TGM2) changes represent the potential for extracellular matrix restructuring and immune cell infiltration similar to MI.^42^ Significant changes in CR2 and CR1 at week 16 post-PAB suggest a complement-induced damage response for immune cell infiltration in remodeling similar to MI.^43^

One of the most logical intervention genes from our data is *FLNA*. Multiple transcripts were altered due to PAB for Filamin A (*FLNA*), which was significantly lower in both MV and TV. The sex (X)-chromosome encoded FLNA cytoskeletal protein is involved in the organization of mature leaflets through GTPase-mediated signal transduction,^44^ with clinical loss-of-function and missense variants resulting in a range of disorders that include tricuspid aortic valve disease and PH.^45,46^ FLNA knockdown protects against hypoxia-induced damage and remodeling during PH through small GTPase signaling.^47^ Given that decreased expression of *FLNA* is found at 16 weeks and not 8 weeks, it suggests that the change in expression correlates to the remodeling of the TV. FLNA has a known role in endocrine tumors through various endocrine substrates, such as IGF2.^48^ Advances in identifying *FLNA* as a master regulator of cardiovascular remodeling^49^ have increased therapeutic studies for FLNA targeting.^50^ Further investigation into how these transcriptional alterations might affect the otherwise unstressed MV in this model may provide insights into the ramifications of PH-induced TR and the role of endocrine signaling throughout the heart between weeks 8 and 16 post-PAB.

## Conclusions

Treating PH, RHF, and TR remains challenging, with diagnosis, prevention, and treatment strategies still requiring more development. Advancing techniques and technologies offer the ability to comprehend the complex interaction between physiological changes and gene expression. Our results support the role of delayed TV transcriptional changes in PAB. This work supports the need to expand human TV tissue collection followed by omic investigations into changes. Our data suggests that the TV may be a potential target for therapeutic interventions rather than just mechanical alterations. There is a need for expanded technology integrations to improve our ability to understand gene regulation and cellular changes, such as long-read RNA sequencing, single-cell profiles, and spatial transcriptomics in both the sheep PAB model and in human tissues. A complete understanding of valve dynamics and their interaction with disease states will require a focus on all heart regions to further dissect the temporal changes, with the potential for the identified endocrine signals to give rise to therapeutic interventions.

## Acknowledgments

This work was supported by the Alliance Foundation of Michigan State University and Corewell Health (to AA and TAT), Corewell Health Heart and Vascular Institute, American Heart Association grant 23SCISA1144887 (to JWP), National Institutes of Health 5R01AI171984 (to JWP) and R01HL165251 (to MKR and TAT).

**Footnote**

## Nonstandard Abbreviations and Acronyms

RHF: right-sided heart failure
LHF: left-sided heart failure
TR: tricuspid valve regurgitation
PH: pulmonary hypertension
PAB: pulmonary artery banding
TPM: transcript per million
PCA: principal component analysis
OMIM: online mendelian inheritance in man
OTP: open target platform
IMPC: international mouse phenotyping consortium
MV: mitral valve
AML: anterior mitral valve leaflet
PML: posterior mitral valve leaflet
TV: tricuspid valve
ATL: anterior tricuspid valve leaflet
PTL: posterior tricuspid valve leaflet
STL: septal tricuspid valve leaflet
LV: left ventricle
RV: right ventricle
PA: pulmonary artery
MPAP: mean pulmonary artery pressure
RAarea: right atrium area
RVd1: right ventricle diameter
TAPSE: tricuspid annular plane systolic excursion
RVFAC: right ventricle fractional area contraction

## Notes

### Competing Interest Statement

The authors have declared no competing interest.

